# Evaluation of a high-throughput deorphanization strategy to identify cytochrome p450s important for odor degradation in *Drosophila*

**DOI:** 10.1101/2021.08.06.455295

**Authors:** Shane R. Baldwin, Pratyajit Mohapatra, Monica Nagalla, Rhea Sindvani, Desiree Amaya, Hope A. Dickson, Karen Menuz

## Abstract

Members of the cytochrome p450 (CYP) enzyme family are abundantly expressed in insect olfactory tissues, where they are thought to act as Odorant Degrading Enzymes (ODEs). However, their contribution to olfactory signaling *in vivo* is poorly understood. This is due in part to the challenge of identifying which of the dozens of antennal-expressed CYPs might inactivate a given odorant. Here, we tested a high-throughput deorphanization strategy in *Drosophila* to identify CYPs that are transcriptionally induced by exposure to a plant volatile. We discovered three CYPs selectively upregulated by the odorant using transcriptional profiling. Although these CYPs are shown to be broadly expressed in the antenna in non-neuronal cells, electrophysiological recordings from CYP mutants did not reveal any changes in olfactory neuron responses to the odorant. Neurons were desensitized by pre-exposing flies to the odorant, but this effect was similar in CYP mutants. Together, our data suggest that this transcriptomic approach may not be useful for identifying CYPs that contribute to olfactory signaling. We go on to show that some CYPs have highly restricted expression patterns in the antenna, and suggest that such CYPs may be useful candidates for further studies on olfactory CYP function.

## Introduction

Odors fluctuate rapidly in the environment, and olfactory receptor neurons are able to respond to this dynamic information. Odorants dissolve into the fluid surrounding olfactory receptor neurons, where they bind and activate receptors on the neuron dendrites. It has long been observed that olfactory tissues abundantly express a large variety of xenobiotic metabolizing enzymes, such as esterases, cytochrome p450s, aldehyde oxidases, and glutathione-S-transferases ^1-3^. More than 30 years ago, it was first postulated that these enzymes may contribute to olfactory signaling by terminating the olfactory signal through the conversion of odorants into inactive metabolites ^3^. Since then, studies in humans, rodents, and insects have shown that broad pharmacological blockers of such Odorant Degrading Enzymes (ODEs) can affect the responses of olfactory receptor neurons to odorants ^4-6^. However, there are still relatively few examples of specific ODE genes that are known to affect neuronal responses *in vivo*. Our best understanding of the olfactory function of ODEs is in insects, where several ODEs that metabolize pheromones have been identified and studied *in vitro*.

Cytochrome p450s (CYPs) are found in nearly all organisms and constitute one of the largest gene families ^7^. Members of this large and rapidly evolving enzyme family carry out diverse functions on endogenous and foreign substrates, including hormone synthesis and xenobiotic detoxification. In Drosophila, there are 87 CYPs genes, which are expressed in a variety of tissues ^8-10^. More than 50 CYPs are expressed in the antenna, their primary olfactory tissue ^11,12^. As in other insect species, expression of some *Drosophila* CYPs is highly enriched in the antenna compared to other tissues, suggesting an olfactory function ^12-14^. However, no *Drosophila* CYPs have been reported to contribute to olfactory signaling. In other insects, pharmacological and biochemical studies indicate a role for CYPs in degrading particular odorants and pheromones *in vitro* ^4,15,16^, but to our knowledge, there is only one example of specific CYP shown to affect olfactory neuron signaling *in vivo* ^17^.

A critical hurdle in the identification of CYPs with olfactory roles is that the substrates of most CYPs in *Drosophila*, the most genetically amenable organism, are unknown ^9,18^. Further, many insect CYPs with roles in xenobiotic metabolism act on multiple substrates ^19^, and thus the identification of one substrate does not preclude another. Additionally, it is not yet possible to predict substrates from CYP sequences alone because they have highly divergent protein sequences, despite their conserved three dimensional structure ^20^. It is estimated that one-third of the *Drosophila* CYP family is involved in metabolizing environmental xenobiotics based on their evolutionary instability, enrichment in tissues commonly associated with detoxification, and ability to be induced by xenobiotics ^18,21^. Of these, ∼90% are expressed in the Drosophila antenna ^11,12^. Given this large number of candidates and the hundreds of odors known to activate *Drosophila* olfactory neurons, it is difficult to predict which CYP may act on which odorant.

In *Drosophila* as in other insects, the presence of xenobiotics leads to the transcriptional upregulation of selective CYPs, a process referred to as induction ^22-26^. Although CYP induction is generally measured following xenobiotic feeding or physical contact, some recent studies in beetles, locusts, and moths have indicated that exposure to volatile odorants can induce CYP expression in the antenna ^27-30^. In this study, we tested whether CYPs can also be induced by odorant exposure in *Drosophila*, and if so, whether the induced CYPs might contribute the detection of that odorant by olfactory neurons.

## Results

### Odorant exposure leads to transcriptional upregulation of a small subset of CYPs

To identify CYPs that may act on a particular odorant, we evaluated transcriptional deorphanization, an approach similar to a strategy employed to deorphanize mammalian odorant receptors ^31^. We exposed flies to a high concentration (5%) of geranyl acetate, ethyl lactate, or the solvent DMSO for five hours. Although the stimulation paradigm is non-physiological, it was used with the goal of strongly driving the upregulation of relevant CYPs. We then harvested three biological replicates of antennal RNA from each condition and used the RNA for RNASeq transcriptional profiling (Fig. 1A).

**Fig. 1:**
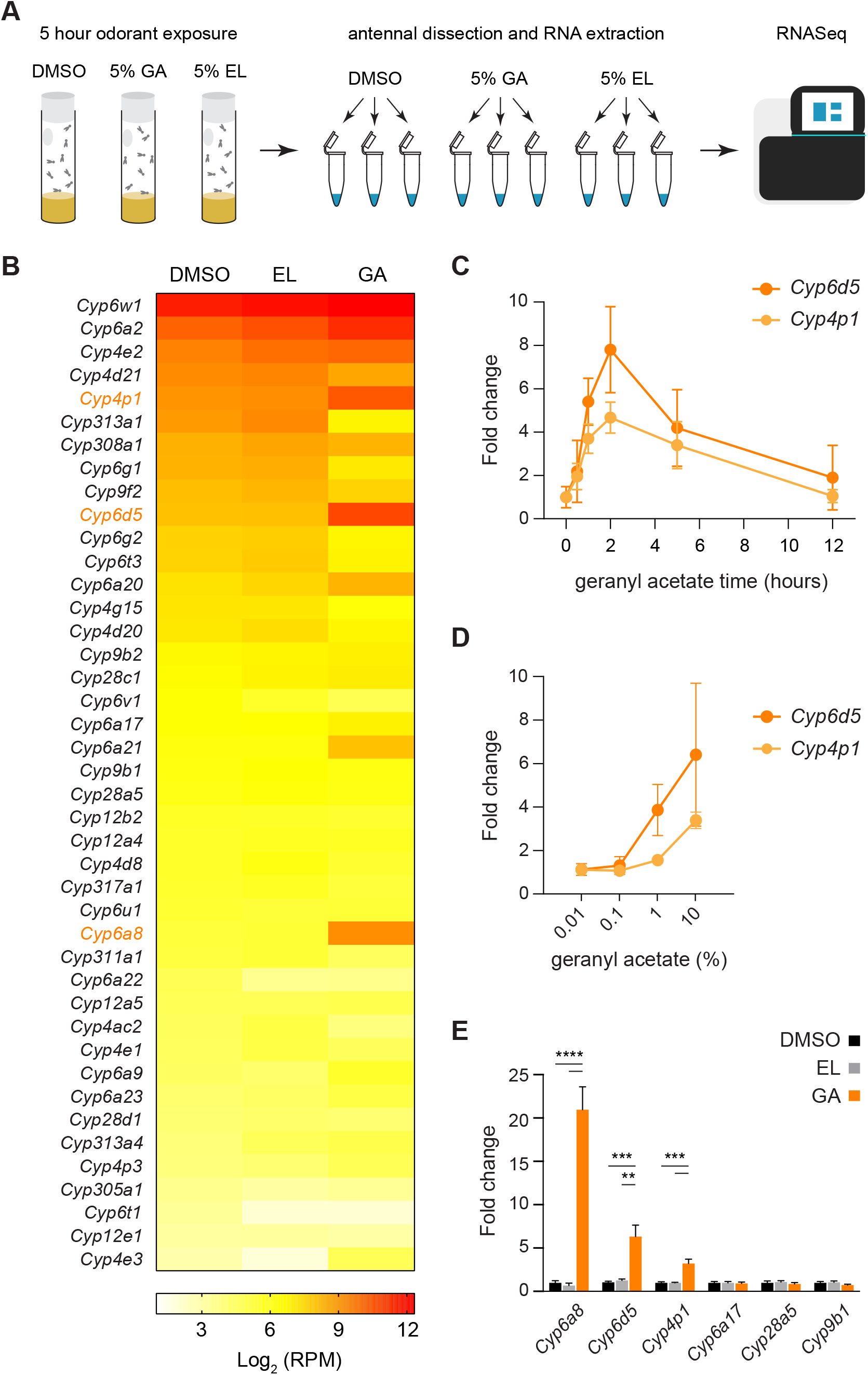
Three CYPs are consistently and specifically upregulated by geranyl acetate exposure. (A) Schematic of RNASeq workflow. Flies were exposed for 5 hours to either DMSO, 5% geranyl acetate (GA), or 5% ethyl lactate (EL). Three biological replicates of ∼200 antenna were collected and used for RNASeq analysis. (B) A heat map showing the average expression in log_2_(RPM) of each of the 42 antennal-expressed CYP genes in each odor exposure condition (n=3). GA-induced CYPs are in orange font. (C) The antennal expression of *Cyp6d5* and *Cyp4p1* detected with qRT-PCR following exposure to 5% geranyl acetate for varying lengths of time. Expression is normalized to expression at 0 hours of exposure for each gene (n=4). (D) The antennal expression of *Cyp6d5* and *Cyp4p1* detected with qRT-PCR following 2 hours of exposure to varying concentrations of geranyl acetate in DMSO. Expression is normalized to DMSO exposure alone (n=5-6). (E) The expression of six CYPs following 2-hour exposure to 5% geranyl acetate and measured with qRT-PCR (n=5). Expression is normalized to DMSO exposure alone. Data were analyzed by one-way ANOVAs followed by Tukey post-hoc tests. Statistical significance is presented as **p<0.01, ***p<0.001, and ****p<0.0001. Other comparisons were not significant (p>0.05). For C-E, data are shown as the mean ± SEM.

The expression level of most CYPs was stable in the presence of either odorant (Fig. 1B and Supplementary Table S1). Using the EdgeR program for differential gene expression analysis, we identified three CYPs, *Cyp6a8, Cyp6d5*, and *Cyp4p1*, that were induced by geranyl acetate exposure compared to either DMSO or ethyl lactate exposure with FDR<0.15 (Supplementary Table S1). No CYPs were induced by ethyl lactate. *Cyp6a8* showed the highest fold-change (∼15-fold) following exposure to geranyl acetate, and it had the lowest DMSO expression (37 RPM) of the geranyl-acetate sensitive CYPs. *Cyp4p1* had the smallest fold-change (∼2.7-fold), and the highest baseline expression (486 RPM), whereas *Cyp6d5* was intermediate in both measures.

We next measured the timecourse of antennal CYP induction following geranyl acetate exposure using qRT-PCR. We found that transcript levels of *Cyp4p1* and *Cyp6d5* rose sharply during the first two hours of geranyl acetate exposure, and then fell gradually over the next 10 hours (Fig. 1C). This is in agreement with a timecourse previously observed for CYP induction in *Drosophila* ^24^. Given the peak response at two hours, further experiments were carried out at this time point. Accurate detection of *Cyp6a8* expression was difficult due its relatively high C_T_ value reflecting its low baseline expression, and it was not analyzed for these experiments.

We next examined the dose-response relationship for CYP induction by geranyl acetate exposure. We found that >0.1% geranyl acetate was needed to elicit changes in *Cyp6d5* or *Cyp4p1* expression in the antenna, and that their upregulation did not saturate even at 10% geranyl acetate (Fig. 1D).

Finally, we validated the selective induction of *Cyp4p1, Cyp6d5*, and *Cyp6a8* by geranyl acetate by exposing flies to 5% geranyl acetate, ethyl lactate, or DMSO for two hours and quantifying antennal gene expression with qRT-PCR. Consistent with our RNASeq data (Fig. 1B), *Cyp4p1, Cyp6d5*, and *Cyp6a8* were consistently upregulated by geranyl acetate exposure, but not by exposure to ethyl lactate (Fig. 1E). In contrast, expression levels of three control CYPs, *Cyp6a17, Cyp28a5* and *Cyp9b1*, were unchanged following either geranyl acetate or ethyl lactate exposure. Thus, geranyl acetate induces a selective upregulation of *Cyp4p1, Cyp6d5*, and *Cyp6a8*, which we will refer to as GA-induced Cyps.

### *Cyp4p1* and *Cyp6d5* are broadly expressed in non-neuronal cells

We investigated the localization of GA-induced CYPs in antennal sections using fluorescence *in situ* hybridization (FISH). Anti-sense FISH probes detected specific expression of *Cyp4p1* and *Cyp6d5*, but not *Cyp6a8*, the CYP with lowest antennal expression (Fig. 1B and Supplementary Fig. S1). Staining antennal sections from flies exposed to geranyl acetate revealed that both *Cyp4p1* and *Cyp6d5* are broadly expressed throughout the antenna (Fig. 2A,B). Given that neurons responding to geranyl acetate are only found in one of ∼20 classes of olfactory sensilla ^32-35^, this suggests that these CYPs are not exclusively expressed in geranyl acetate sensitive sensilla.

**Fig. 2:**
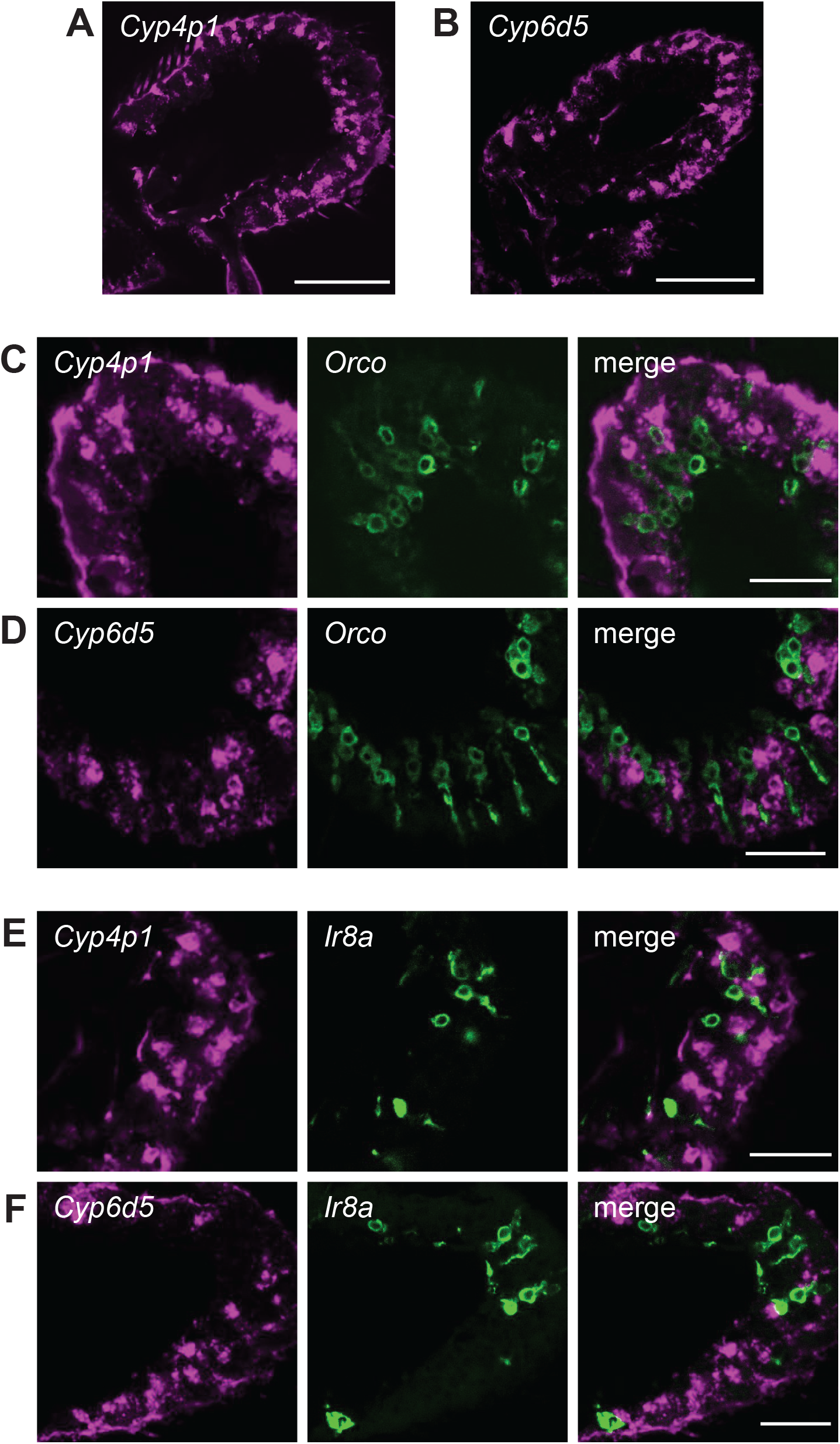
*Cyp4p1* and *Cyp6d5* are broadly expressed and are not found in olfactory neurons. (A, B) Antennal sections from flies exposed to geranyl acetate for two hours and labeled with fluorescence *in situ* hybridization (FISH) probes (magenta) for (A) *Cyp4p1* or (B)*Cyp6d5*. Scale bars, 50 µm. (C, D) Antennal sections from geranyl-acetate exposed *Orco*>*GFP* flies labeled with an anti-GFP antibody (green) and FISH probes (magenta) for (C) *Cyp4p1* or (D) *Cyp6d5*. Scale bars, 20 µm. (E, F) Similar to (C, D), but with *Ir8a*>*GFP* flies.

Within the antenna, auxiliary cells are thought to express Odorant Degrading Enzymes (ODEs), including CYPs, but few studies have examined their cellular localization. In mammals, ODEs have been detected in both olfactory neurons and surrounding support cells ^3^. Might some CYPs likewise be expressed in olfactory neurons in addition to their expected expression in auxiliary cells? To address this question, we examined *Cyp4p1* and *Cyp6d5* expression with FISH on antennal sections from flies in which subsets of olfactory neurons were labeled with GFP driven by transgenic reporters. In *Drosophila*, >70% of olfactory neurons express Orco, a co-receptor required for the function of OR type olfactory receptors ^36,37^. We found that Orco^+^ neurons do not co-express either *Cyp4p1* or *Cyp6d5* (Fig. 2C,D). An IR-family co-receptor, Ir8a, is expressed by nearly olfactory neurons that do not express Orco ^36^. Like Orco^+^ neurons, Ir8a^+^ neurons express neither *Cyp4p1* nor *Cyp6d5* (Fig. 2E,F). Taken together, our data indicate that the GA-induced CYPs are not expressed in olfactory neurons.

### Loss of GA-induced CYPs does not affect electrophysiological responses of olfactory neurons

Do *Cyp4p1, Cyp6a8*, and *Cyp6d5* contribute to olfactory responses mediated by geranyl acetate sensitive olfactory neurons? We hypothesized that if these CYPs inactivate geranyl acetate, their absence might prolong or amplify neuronal spiking responses to this odor. To test this possibility, we generated mutants for each of the three GA-induced CYPs using CRISPR/Cas9 engineering to replace much of the coding region of each gene with a 3xP3-DsRed reporter (Methods). The conserved CYP consensus sequence containing a required cysteine that interacts with the heme iron was deleted in each mutant, presumably rendering each CYP non-functional ^9,38^. Following PCR validation, the deletions were outcrossed for at least 10 generations to a standardized genetic background. The *Cyp4p1*^*1*^ and *Cyp6a8*^*1*^ mutants were viable, but we were unable to obtain *Cyp6d5*^*1*^ homozygous mutants.

We used single-sensillum recordings (SSR) to examine the spiking responses of geranyl acetate-sensitive olfactory neurons in the *Cyp4p1*^*1*^ and *Cyp6a8*^*1*^ flies. In *Drosophila*, only Or82a^+^ neurons in ab5 sensilla respond strongly to geranyl acetate ^32-35^. We found that responses to brief (500 ms) pulses of geranyl acetate were similar in the two CYP mutants as in the genetic background control (Fig. 3A). Likewise, responses to another odorant detected by Or82a, citral, were unaffected by the loss of either *Cyp4p1* or *Cyp6a8* (Fig. 3A). We next investigated spiking responses to geranyl acetate using a dose-response curve, and found that the response EC_50_ in control flies was not statistically different from either CYP mutant (Fig. 3B).

**Fig. 3:**
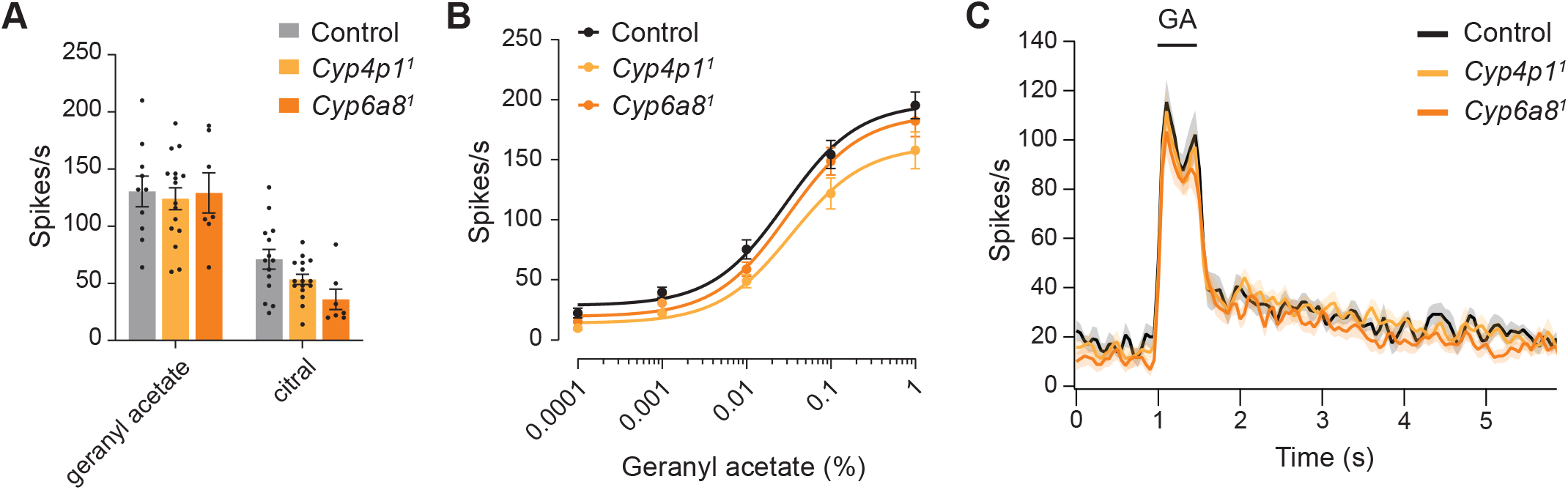
Odor responses of GA-sensitive neurons are unaffected in *Cyp4p1*^*1*^ and *Cyp6a8*^*1*^ mutants. (A) Responses to odors that activate the ab5A neuron, geranyl acetate (0.1%) and citral (0.1%), are similar in *Cyp4p1*^*1*^ and *Cyp6a8*^*1*^ mutants as in controls. Bar graphs depict the mean ± SEM overlaid with the individual data points. Data analyzed with two-way ANOVA followed by Tukey’s post-hoc test (n=8-17 sensilla each, p>0.05 for all comparisons). (B) Dose-response curve of spiking responses to geranyl acetate. Dose response curves show the mean ± SEM and the curve fit to the Hill equation. The EC_50_s were not significantly different between the three genotypes (control 0.029% ± 0.008, *Cyp4p1*^*1*^ 0.034% ± 0.013%, *Cyp6a8*^*1*^ 0.031% ± 0.010%, one-way ANOVA, n=8-15 sensilla each). (C) Peristimulus time histograms (PSTHs, mean ± SEM) show spiking responses to a 500 ms pulse of geranyl acetate (0.01%). Each trace (colored line) represents the mean of 8-15 recordings taken from the data for the dose-response curve in 3C.

Data in Fig. 3A and 3B quantified the cumulative spiking response over the 500ms response period. However, this could obscure more subtle changes in the response kinetics. We therefore inspected the instantaneous firing frequency over a longer time period using peristimulus time histograms. The kinetics of the geranyl acetate responses were indistinguishable from controls in the *Cyp4p1*^*1*^ and *Cyp6a8*^*1*^ mutants (Fig. 3C).

We next examined responses to repetitive stimuli, for which odorant inactivation may be more critical compared to responses to single pulses. In the absence of odorant removal, modeling suggests that odorants accumulate in the sensillum lymph and would not reflect environmental levels ^39^. We therefore assessed spiking responses to paired-pulses and trains of geranyl-acetate stimuli in *Cyp4p1*^*1*^ mutants. First, we applied two 500 ms pulses of geranyl acetate at 10, 3, or 1.5 second intervals, and we measured the paired-pulse ratio for each sensillum as the response to the second pulse divided by the first pulse. At each interstimulus interval (ISI), the paired-pulse ratio was ∼1 in both control and *Cyp4p1*^*1*^ flies, and their ratios were not statistically different from each other (Fig. 4A-D).

**Fig. 4:**
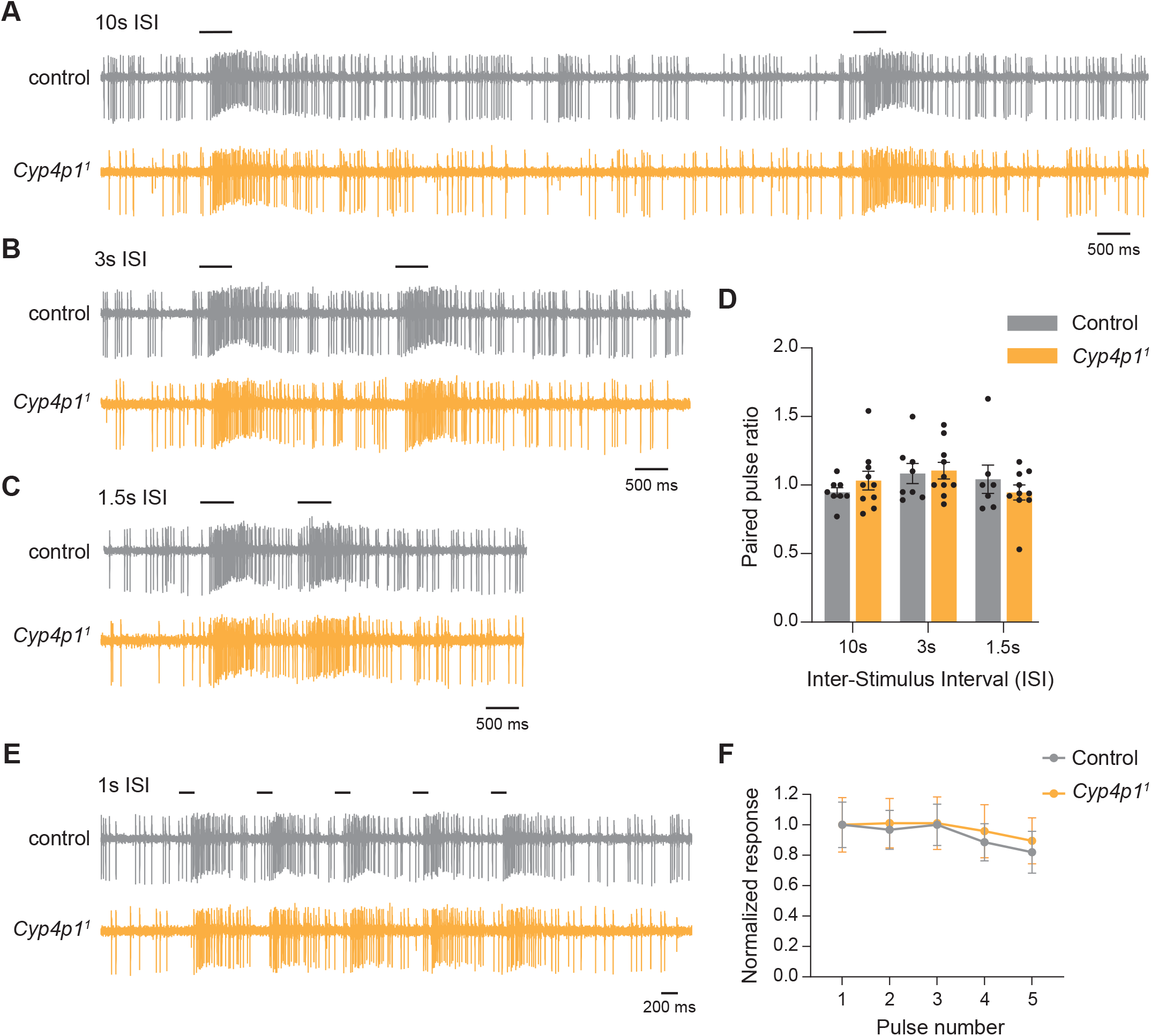
Responses to repetitive geranyl acetate pulses are unaffected in *Cyp4p1*^*1*^ mutants. (A) SSR traces from control (gray) and *Cyp4p1*^*1*^ (orange) flies showing responses to paired 500 ms pulses of geranyl acetate (0.01%) given 10 seconds apart. The time at which pulses were given is shown with a black bar. (B, C) Similar to A, but for with interstimulus intervals (ISI) of (B) 3 seconds or (C) 1.5 seconds. (D) Quantification of the paired-pulse ratio at each ISI. Bar graphs depict the mean ± SEM overlaid with the individual data points. The ratios were similar in control and *Cyp4p1*^*1*^ flies (two-way ANOVA, p> 0.05, n=7-10 each). (E) SSR traces from control (gray) and *Cyp4p1*^*1*^ (orange) flies showing responses to a train of five 200 ms pulses of geranyl acetate (0.01%) given 1 second apart. (F) Spiking responses to the geranyl acetate train as a function of the pulse number. Responses for each sensillum were normalized to the mean response to the first pulse for that genotype. The graph shows the mean ± SEM. Responses slightly decrease with increasing pulse number. The responses were similar in control and *Cyp4p1*^*1*^ flies (two-way repeated measures ANOVA, p> 0.05, n=5 and 7, respectively).

We also applied a train of five 200 ms geranyl acetate pulses at 1 Hz, and measured the response to each pulse relative to the response to the first pulse for that sensillum. Although responses trended slightly lower with increasing pulse number, no changes were found in the response of *Cyp4p1*^*1*^ mutants relative to controls (Fig. 4E,F).

### Reduced spiking responses to geranyl acetate following odorant exposure are independent of GA-induced CYPs

Although our data do not reveal a role for GA-induced CYPs under baseline conditions, we wondered whether they might play a greater role following their upregulation by geranyl acetate exposure. For example, the tonic presence of geranyl acetate might desensitize Or82a receptors, reducing its odorant sensitivity, and the absence of odorant clearance might exacerbate this effect. To test this possibility, we pre-exposed flies to 5% geranyl acetate for two hours and examined ab5 spiking responses using electrophysiological recordings within 2 hours following the odor exposure. Strikingly, responses to both 0.01% and 0.1% geranyl acetate in control flies were reduced ∼70% by the prolonged exposure to geranyl acetate (Fig. 5A,B).

**Fig. 5:**
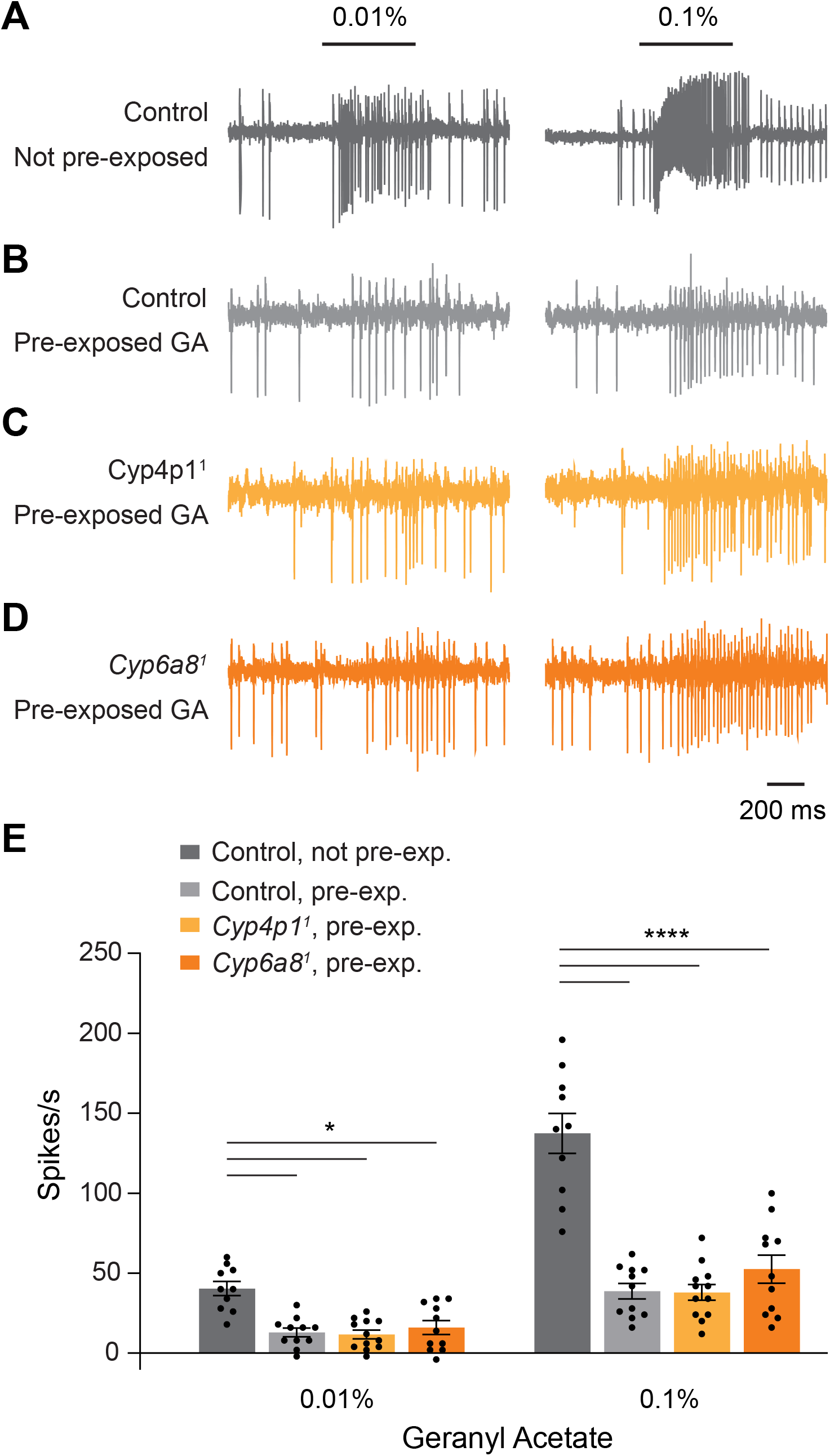
Pre-exposure to geranyl acetate reduces spiking responses to geranyl acetate. (A) SSR traces show responses of an ab5 sensillum to a 500 ms pulse of 0.01% and 0.1% geranyl acetate in control flies not exposed to geranyl acetate. (B) SSR traces show responses of an ab5 sensillum to geranyl acetate in control flies following two hours of exposure to 5% geranyl acetate. (C,D) Similar to B, but for (C) *Cyp4p1*^*1*^ and (D) *Cyp6a8*^*1*^ mutants. (E) Quantification of spiking responses to geranyl acetate. Bar graphs depict the mean ± SEM overlaid with the individual data points. Data were analyzed with a two-way ANOVA followed by Tukey’s post-hoc test. Statistical significance is presented as *p<0.05 and ****p<0.0001. Other comparisons were not significant (p>0.05).

We also examined responses in *Cyp4p1*^*1*^ and *Cyp6a8*^*1*^ flies pre-exposed to geranyl acetate. They exhibited suppressed spiking responses to geranyl acetate, similar to controls (Fig. 5B-E). Therefore, the mechanism by which odorant exposure leads to reduced responses is independent of the expression of these two CYPs.

### An alternative strategy to identify CYPs with roles in *Drosophila* olfaction?

Together, our data indicate that although the expression of some CYPs is induced by geranyl acetate, they are unlikely to play a role in olfactory signaling mediated by this odorant. This suggests that transcriptional deorphanization may not be a useful approach to identify CYPs with roles in *Drosophila* olfaction. We wondered whether investigating the expression pattern of antennal CYPs may be a useful strategy to explore this question in the future. Although the GA-induced CYPs we identified are broadly expressed, there may be some Cyps whose expression is limited to particular sensillar classes, and thus may contribute to olfactory signaling by the neurons housed in those sensilla.

Surprisingly few studies have investigated the localization of CYPs within antennae in either *Drosophila* or other insects ^17,40^. To explore whether CYPs with restricted expression exist in *Drosophila*, we selected four antennal CYPs for reporter line generation: 1) *Cyp4e2*, one of the most highly expressed CYPs in the antenna ^11^, 2) *Cyp308a1*, a CYP exclusively found in the antenna ^12,13^, and 3) *Cyp313a4* and *Cyp4d21*, whose expression is greatly reduced in *atonal* mutants which fail to develop coeloconic sensilla, the sacculus and the arista ^11,41^. Examination of mtdTomato driven by the CYP reporter lines in the antenna revealed that *Cyp308a1* and *Cyp4e2* are widely expressed (Fig. 6A,B). In contrast, expression of *Cyp313a4* is restricted to a small scattered subset of sensilla (Fig. 6C) and *Cyp4d21* expression is limited to the arista (Fig. 6D). We suggest that investigation of *Cyp313a4* and other Cyps with restricted sensillar expression may prove useful in future studies on the role of CYPs in olfactory neuron function.

**Fig. 6.**
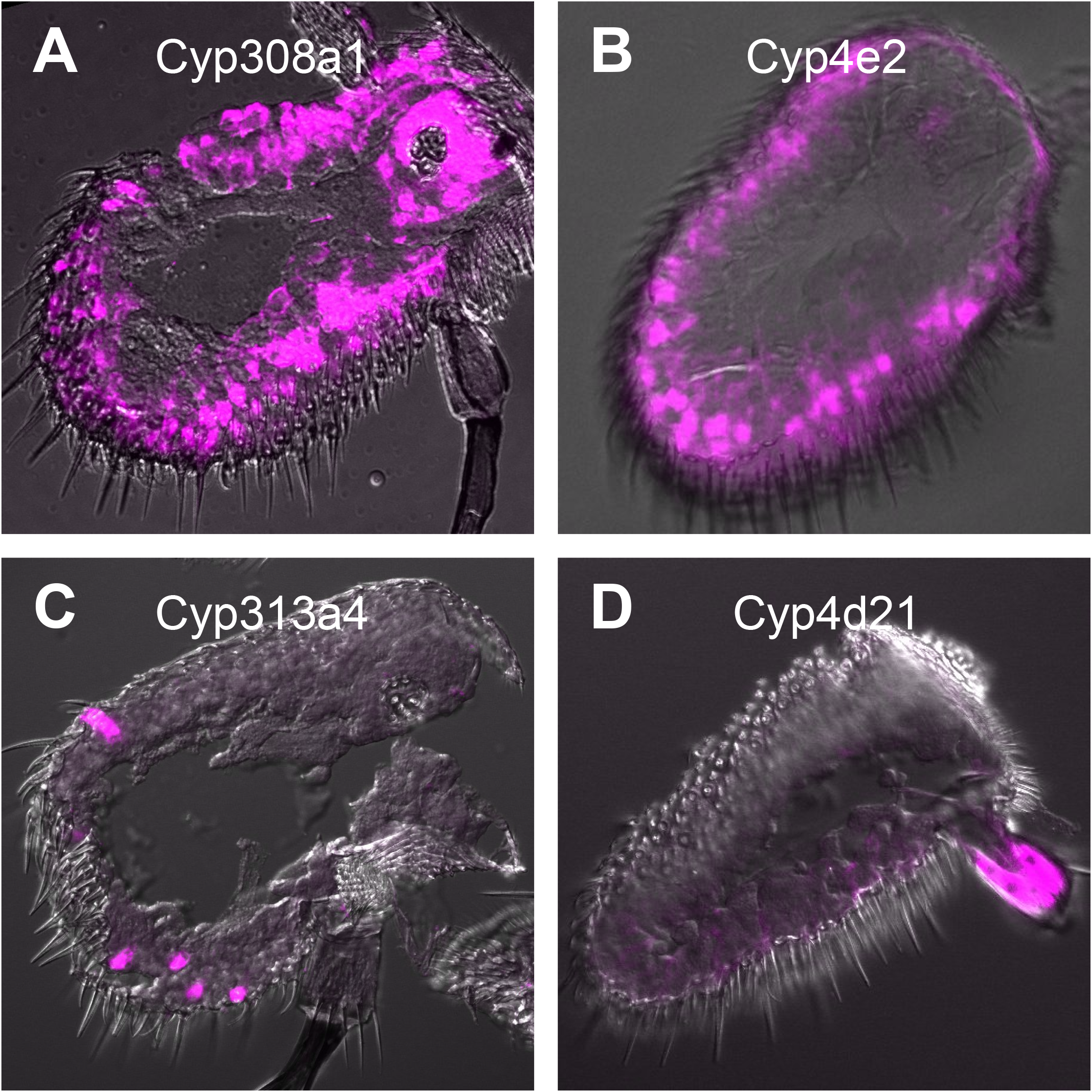
CYPs can have both broad and narrow antennal expression patterns. (A-D) Confocal images of antennal sections labeled with an antibody against mtdTomato (magenta) driven by (A) *Cyp308a1-LexA*, (B) *Cyp4e2-LexA*, (C) *Cyp313a4-LexA*, or (D) *Cyp4d21-LexA. Cyp308a1* and *Cyp4e2* are broadly expressed, whereas *Cyp313a4* is expressed in a subset of sensilla and *Cyp4d21* is expressed in the arista.

## Discussion

In this study, we found that exposure to the odor geranyl acetate specifically and consistently lead to the induction of three CYPs, *Cyp4p1, Cyp6a8*, and *Cyp6d5*, in the antenna. Interestingly, each of these CYPs has been previously shown to be inducible by multiple xenobiotics ^22-25^. The xenobiotics that upregulate these CYPs vary widely in structure. For example, exposure to phenobarbital, caffeine, atrazine, pyrethrum, paraquat, or piperonyl butoxide lead to the induction of one or more of these CYPs ^22,25^. Additionally, each of these three CYPs has a high level of genomic instability, with frequent gene gain and loss events across *Drosophila* species ^21^. These findings, together with the enrichment of *Cyp4p1, Cyp6a8*, and *Cyp6d5* in tissues associated with detoxification such as the midgut, Malpighian tubules and fat body, suggests they function in xenobiotic detoxification rather than the synthesis or degradation of products as part of developmental or physiological processes ^18^. This would be consistent with a role for these CYPs in the degradation of odorants such as geranyl acetate.

We also investigated the localization and function of geranyl acetate-induced CYPs in the antenna. Although bulk RNASeq studies demonstrate that many CYPs are expressed in the *Drosophila* antenna, their localization within this tissue has rarely been examined. Using FISH, we found that *Cyp4p1* and *Cyp6d5* are broadly expressed within the antenna, suggesting they are found in multiple sensilla classes. Further, our data revealed these CYPs are not expressed by olfactory neurons. We suspect that they are expressed by the non-neuronal auxiliary cells, which have long been thought to mediate odorant degradation ^2^.

We then generated homozygous mutants for *Cyp4p1* and *Cyp6a8*, and used these for electrophysiological recordings on the geranyl acetate sensitive ab5 sensilla. We probed multiple aspects of geranyl acetate responsiveness, including sensitivity, kinetics, and repetitive stimuli, but did not find any differences between controls and the CYP mutants. We also examined olfactory neuron responses following prolonged exposure (2 hours) to geranyl acetate, a situation in which the absence of CYPs may enhance odorant accumulation in the sensillum lymph and lead to desensitization. Following geranyl acetate exposure, electrophysiological responses to geranyl acetate were strongly reduced. This is similar to what has been reported for other odorants and is mediated by de-phosphorylation of the Orco co-receptor ^42,43^. However, slow desensitization was not affected by the loss of *Cyp4p1* or *Cyp6a8*.

Our negative results may indicate that odorant degradation mediated by CYPs does not affect olfactory signaling. However, there are caveats that prevent this conclusion. First and foremost, it is unclear whether the induction of a CYP by a xenobiotic compound necessarily implies that the compound is metabolized by that CYP. Although numerous studies demonstrate *Cyp4p1, Cyp6a8*, or *Cyp6d5* induction in *Drosophila*, to our knowledge only one has directly demonstrated that the xenobiotic is modified or degraded by the induced CYP ^25^. More generally, the signal transduction pathway by which xenobiotics induce CYP expression is poorly understood in insects ^26^. As a result, it is possible that xenobiotic exposure leads to the activation of transcription factors that increase expression of a broad array of CYPs, not all of which target the xenobiotic. A second caveat is that CYPs may play redundant roles in odorant degradation, such that the absence of one CYP is compensated by another. We tested each mutant alone, but not in combination. Finally, it is possible that *Cyp6d5* acts on geranyl acetate, but the lethality of our *Cyp6d5*^*1*^ mutant prevented us from testing the role of this enzyme. Given these caveats, we do not believe our data can rule out a role for CYPs in olfactory signaling.

Given the limitations of a transcriptional deorphanization approach, how might the role of CYPs be studied in *Drosophila* olfaction in the future? We have begun to generate a collection of transgenic CYP reporter lines to investigate their expression patterns within the antenna. We have identified some CYPs with highly restricted expression patterns, such as *Cyp313a4*. Such CYPs most likely act on a narrow range of substrates rather than fulfilling a general metabolic role, and quite likely these substrates would include odorants detected by that sensillum. We propose that identifying the sensilla classes in which CYPs with restricted expression are found and then testing their role in responses mediated by neurons in those sensilla will be informative in the future.

## Methods

### Fly lines

Drosophila melanogaster flies were raised on a standard cornmeal molasses food at 25°C in an incubator with a 12:12 hour day/night cycle. *Canton-S* flies were used for RNA sequencing and qRT-PCR. FISH was carried out on lines in which either *Ir8a-Gal4* ^11^ or *Orco-Gal4* ^44^ drove expression of *UAS-mCD8::GFP* ^45^. LexA lines for *Cyp308a1, Cyp4e2, Cyp313a4*, and *Cyp4d21* were generated as described below and drove expression of LexAop-mtdTomato ^46^. *Cyp4p1*^*1*^ and *Cyp6a8*^*1*^ loss-of-function mutations, generated as described below, were outcrossed for 10 generations to a white-eyed *Canton-S* (*wCS*) line prior to electrophysiological recordings ^46^. The *wCS* line served as a control for all electrophysiological experiments.

### Generation of transgenic reporter lines

Transgenic LexA reporter lines for *Cyp4d21, Cyp308a1, Cyp4e2*, and *Cyp313a4* were generated using standard methods, similar to described previously ^11,46^. The 5’ and 3’ regions flanked the genes were cloned from the following tiling bacterial artificial chromosomes (BACs) corresponding to the *Drosophila melanogaster* reference genome: BACR13D17 (*Cyp4d21*), BACR07J06 (*Cyp313a4*), BACR14D22 (*Cyp308a1*), and CH322-97G21 (*Cyp4e2*), ^47,48^. Multisite Gateway Pro Recombination (Thermo Fisher) was used to assemble the 5’ and 3’ genomic regions surrounding *LexA* in the pBGRY destination vector ^46^. For *Cyp4e2-LexA*, the 5’ region included chromosome 2R: 8,442,589-8,444,770 and the 3’ region 2R: 8,446,895-8,447,622. For *Cyp4e2-LexA*, the 5’ region included chromosome 2L: 7,597,215-7,605,812 and the 3’ region 2L: 7,607,235- 7,610,142. For *Cyp313a4-LexA*, the 5’ region included chromosome 3R: 12,681,257-12,683,325 and the 3’ region 2L: 12,678,913-12,679,412. For *Cyp308a1-LexA*, the 5’ region included chromosome X: 18,705,774-18,709,894 and the 3’ region X: 18,711,500-18,716,181. Plasmids were verified with restrictions digests and sequencing. The vectors were injected *Drosophila* embryos by BestGene, Inc. for PhiC31-mediated genomic integration ^49^. Vectors were integrated into either attP40 (2nd chromosome, *Cyp4e2-LexA, Cyp4d21-LexA*, and *Cyp308a1*), VK27 attP-9A (3^rd^ chromosome, *Cyp4e2-LexA* and *Cyp4d21-LexA*), or attP2 (3^rd^ chromosome, *Cyp313a4-LexA*) landing sites ^50-52^. Flies with successful integration were identified by expression of *mini-white* in their eyes.

### Generation of CYP mutants

*Cyp4p1*^*1*^, *Cyp6a8*^*1*^, and *Cyp6d5*^*1*^ mutant fly lines were generated with CRISPR/Cas9 engineering and homology directed repair, similar to described previously ^53^. Of the coding sequences for each gene, 94% are deleted in *Cyp4p1*^*1*^, 35% for *Cyp6a8*^*1*^, and 54% for *Cyp6d5*^*1*^. For each gene, guide RNA (gRNA) sequences were inserted into pU6-BbsI-chiRNA ^54^ using the Q5 site-Directed Mutagenesis Kit (New England Biolabs). Each PCR used a common primer GAAGTATTGAGGAAAACATA and a primer containing the gene specific gRNA sequence followed by GTTTTAGAGCTAGAAATAGC. Correct guide-RNA plasmids were verified by sequencing. For each gene one or two gRNA sequences were generated with the following primers: GCCAGTAGAGCAGGGCACTT and GCAGAAGATTGTCTTCCATA for *Cyp4p1*, GGTTAGCTCGTATAGGGTAA for *Cyp6d5*, and GAAAGACGGAATCTATCTGA and GATTCTTTGCCAGTTCGTAT for *Cyp6a8*. Donor plasmids for homology directed repair were generated by cloning homology arms from *Canton-S* genomic DNA with Phusion Hot Start DNA polymerase (New England Biolabs) and inserting them sequentially into the pHD-DsRed-attP vector using restriction enzymes ^55^. The homology arms flanked the 3xP3-DsRed marker after being inserted into the vector. The homology arms for *Cyp4p1* spanned from chromosome 2R: 9,239,359-9,240,112 and 2R: 9,241,981-9,242,939. The homology arms for *Cyp6a8* spanned from chromosome 2R: 14,886,540-14,887,420 and 2R: 14,888,010-14,888,943. The homology arms for *Cyp6d5* spanned from chromosome 3R: 14,028,829-14,029,440 and 3R: 14,030,647-14,031,375. Donor plasmids and gRNA plasmids were injected into embryos by BestGene, Inc. The injected embryo lines were *y,w*;;*nos-Cas9*(III-attP2) for Cyp4p1 and Cyp6a8 and *y,w*;*nos-Cas9*(II-attP40) for Cyp6d5 ^56^.

Ocular DsRed expression was used to identify flies in which the donor plasmid had been integrated into the genome. Sequencing of PCR products was used to validate the expected genetic changes. Homozygous flies were obtained for mutations in *Cyp4p1* and *Cyp6a8* mutations, but not for *Cyp6d5*. Flies were outcrossed for 10 generations to the *wCS* genetic background prior to experiments.

### Antennal immunohistochemistry

Flies aged approximately one week were anesthetized, aligned in a collar, and encased in OCT (Tissue-Tek) up to their necks in a silicone mold. Following freezing on dry ice, heads were snapped off and stored as blocks at −80°C. Sections (20 µm) were collected on a Leica cryostat and used for immunohistochemistry. Slides were fixed in 4% formaldehyde in PBS for 10 minutes, washed three times in PBS, permeabilized in PBS with 0.1% Tween (PBST) for 30 minutes, and then immersed in blocking buffer (PBST with 1% BSA) for 30 minutes. Sections were incubated overnight at 4°C in rabbit anti-RFP antibody (Rockland, RRID:AB_2209751) diluted 1:500 in blocking buffer. Following removal of excess primary antibody by three washes in PBST, the secondary antibody goat anti-rabbit (Fisher, RRID:AB_143157) was applied at 1:500 diluted in blocking buffer for two hours at room temperature. Sections were then washed three times in PBST and mounted in Vectashield. Imaging was performed with a Nikon A1R confocal microscope at Advanced Light Microscopy Facility at the University of Connecticut. Images were process with ImageJ/FIJI software.

### RNA Sequencing

Canton-S flies (∼100) aged 7-10 days were exposed for 5 hours to 5% geranyl acetate in DMSO, 5% ethyl lactate in DMSO, or 20 µl DMSO. Exposures took place in a 25°C incubator in narrow plastic fly vials containing fresh food and a 13 mm GE Whatman antibiotic disc with 20 µl of the odor solution. Exposure vials were kept in 500 ml screw-top containers to prevent mixing of odors. After 5 hours, vials were placed at −20°C until flies were immobilized. Flies were flash frozen in liquid nitrogen, and the antennae were manually dissected into a 1.5 ml Eppendorf tube set in a liquid nitrogen bath. After collection of ∼200 antennae, the antennae were pelleted by centrifugation at 4°C. Three independent antennal samples were collected for each odor condition. Antennal tissues were ground using an RNAse-free pestle, and total RNA was extracted using QIAShredders (Qiagen) and the QIAGEN RNeasy Micro kit. The RNeasy Micro kit protocol was followed as directed. The RNA was then cleared of genomic DNA using DNAse from an iScript gDNA Clear cDNA Synthesis Kit (Bio-Rad).

The nine RNA samples (∼0.5 µg each) were sent to the Center for Genome Innovation at the University of Connecticut for quality control, library preparation, and sequencing. Libraries were prepared using the Illumina mRNA sample prep kit for non-stranded RNA, and sequencing was carried out on an Illumina NextSeq 500 with a mid output 150 cycle sequencing run to produce paired-end reads.

### RNA Sequencing Analysis

Raw reads were obtained from BaseSpace Sequence Hub (Illumina). Sickle was used with the default parameters to perform quality control and produce trimmed reads ^57^. STAR was used with the default parameters to align trimmed reads to the *Drosophila melanogaster* reference genome (dm6 2014, Release 6 plus ISO1 MT) with mitochondrial genes removed ^58,59^. Raw counts of reads mapping uniquely to each gene were obtained using HTSeq-count with the “intersection-nonempty” and “nonunique-none” modes (Anders et al., 2015). Mapped reads are available at the NCBI SRA under bioProject accession number PRJNA746306.

The Reads per Million mapped reads (RPM) for each gene was calculated by dividing the raw reads by total reads, then multiplying by 1,000,000. The 87 CYPs in the *Drosophila* genome were identified within the dataset, and the 42 CYPs whose expression was >5 RPM in each DMSO-exposed sample were considered to be antennal-expressed (Table S1). An ExactTest was run in EdgeR v3.26.8 ^60^ in R v3.6.1 to identify differentially expressed genes between flies exposed to geranyl acetate or to DMSO. Separate paired ExactTests were run for ethyl lactate and DMSO, and ethyl lactate and geranyl acetate. The 42 antennal-expressed CYPs were extracted from each analysis. Of these, five CYPs were differentially expressed with FDR<0.15 between flies exposed to geranyl acetate and those exposed to DMSO (Table S1). These CYPs were also differentially expressed between flies exposed to geranyl acetate and those exposed to ethyl lactate. Of these, *Cyp6a8, Cyp6d5*, and *Cyp4p1* were upregulated by geranyl acetate exposure, whereas *Cyp313a1* and *Cyp6v1* were downregulated. No CYPs were differentially expressed between flies exposed to ethyl lactate and those exposed to DMSO.

### qRT-PCR analysis

Canton-S flies (∼100) aged 7-10 days were exposed to 20 µl of 5% geranyl acetate in DMSO, 5% ethyl lactate in DMSO, or DMSO, in the same manner as for the RNASeq experiment. Flies were exposed for two hours, except when examining the time course of the transcriptional response when they were exposed for 0 to 12 hours. Odor exposure and total RNA isolation was carried out as described above. Total RNA (280 ng per sample) was used to generate cDNA using the iScript gDNA Clear cDNA Synthesis Kit (Bio-Rad). The cDNA was used as a template for qRT-PCR with SsoFast EvaGreen Supermix (Bio-Rad). A CFX96 thermocycler (Bio-Rad) was used to run the qRT-PCR assays in triplicate. For all experiments, gene expression in each sample was measured relative to the housekeeping gene *eIF1a* to normalize cDNA levels between samples. For the time course analysis, expression at each time point is plotted as the fold change relative to expression at 0 hours of geranyl acetate exposure. For the dose response analysis, expression at each geranyl acetate concentration is plotted as the fold change relative to exposure to DMSO alone. Each reaction was run on 4-6 independent antennal replicates.

### Probe synthesis for fluorescence *in situ* hybridization (FISH)

The *Cyp4p1* gene was amplified from the *Canton-S* antennal cDNA with primers ATGATTATCTTGTGGCTGATTCTG and AATAAGTCACGTTCGCCTCAC and then ligated into pGEM-T Easy (Promega). The vector carrying the full length *Cyp4p1* insert was verified by restriction digests and sequencing. Plasmids containing full length cDNA sequences for *Cyp6d5* and *Cyp6a8* in the pOT2 vector, GH07481 and FI17852 respectively, ^61^, were obtained from the Drosophila Genomics Resource Center.

To generate FISH sense (S) and anti-sense (AS) probes, 1-5 µg of *Cyp4p1*-pGEM, *Cyp6a8*-pOT2 and *Cyp6d5*-pOT2 plasmids were linearized overnight and purified into RNAse-free water using the MinElute Reaction Cleanup Kit (Qiagen). Labeled S and AS probes were generated using the DIG RNA Labeling Kit (SP6/T7) (Roche) per the manufacturer’s instructions. Probes were hydrolyzed for 10 minutes in 30 mM Na_2_CO_3_, 20 mM NaHCO_3_ (pH 10.2). The reaction was terminated with 3M NaOAc, 1% acetic acid (pH 6). Probes were then purified with ethanol precipitation, solubilized and diluted to 50 ng/µL in DEPC water, and stored at −80°C.

### Combined FISH and immunocytochemistry

Groups of 4-7-day old flies were exposed to geranyl acetate or DMSO for two hours. After exposure, flies were anesthetized by cold, maintained on a petri dish on an ice bath, and manually placed on a collar. The heads of 5 males and 5 females were covered with silicone molds containing OCT (Tissue-Tek) and placed on dry ice, and the blocks were snapped off when frozen. The blocks were sectioned at 20 µm using a cryostat. The slides were fixed with 4% paraformaldehyde in PBS for 10 minutes, washed with PBS, and then incubated for 10 minutes in an acetylation solution (0.463g triethanolamine HCl, 56 µL 10N NaOH, 62.5 µL acetic anhydride, 25 mL H20). Slides were then washed in PBS and prehybridized for one hour at 65°C in hybridization buffer (HB: 25 mL formamide, 12.5 ml 20x SSC, 2.5 mg heparin, 0.5 mL 10% Tween-20, 12 mL DEPC H_2_0). DIG labeled probes were diluted to 500 ng/mL in HB, denatured at 80°C for 5 minutes, and cooled on ice for one minute. Slides were covered with 200 µL of probe in HB and a Hybrislip (Sigma-Aldrich). They were then incubated at 65°C in a humid chamber for ∼20 hours. Hybrislips were removed by dunking in 5x SSC warmed to 65°C, and they were then washed three times in 0.2x SSC at 65°C. Slides were then incubated for 10 minutes with TN (100 mM Tris-HCl pH7.4, 150 mM NaCl), and then blocked for 30 minutes in TNB (TN with 1% Blocking Reagent (Roche)) at room temperature. Primary antibody solution containing anti-DIG-POD antibody (1:250, Roche, RRID:AB_514500) and mouse anti-GFP antibody (1:500, Roche, RRID:AB_390913) diluted in TNB was added to each slide under a bridged coverslip and incubated at 4°C overnight. After careful removal of the bridged coverslips, slides were washed in TNT (TN with 0.05% Tween-20). TSA-Cy3 was diluted 1:50 in Amplification Reagent (Perkin-Elmer), and each slide was incubated with 200 µL of this solution in the dark for 10 minutes at room temperature. All further steps were carried out in the dark. Slides were washed in TNT with agitation, and then incubated for 2 hours at room temperature in Alexa Fluor 488 donkey-anti-mouse (1:500 in TNB, Invitrogen, RRID:AB_141607). Following three washes with TNT, and sections were mounted in Vectashield (Vector Labs). Confocal microscopy was performed using a Nikon A1R Confocal microscope in the University of Connecticut Advanced Light Microscopy Facility. All images were processed using FIJI/ImageJ ^62^.

### Odorants

Odorants were geranyl acetate (95%, Alpha Aesar), ethyl lactate (99%, Alpha Aesar), and citral (95%, Sigma). Geranyl acetate and ethyl lactate were diluted into DMSO (Fisher Chemical) for exposing flies prior to RNASeq, qRT-PCR, FISH histology, and electrophysiology. They were used at a 5% concentration, except for the dose-response curve shown in Fig. 1D. For electrophysiological recordings, geranyl acetate and ethyl lactate were diluted 1:10 into paraffin oil (Sigma), and serial dilutions were used to achieve lower concentrations. In Fig. 3A, geranyl acetate was used at 0.1% and citral at 0.1%. In Fig. 3C and 4A-F, geranyl acetate was used at 0.01%.

### Electrophysiology

Single sensillum electrophysiological recordings were performed on 3-7 days old female flies as described ^63,64^. In brief, flies were wedged in a 200 µl pipette tip, exposing a portion of their head and their antennae. One antenna was held gently between a tapered glass electrode and a coverslip. The prep was placed on BX51WI microscope (Olympus) under a continuous 2,000 mL/min humidified air stream delivered through the glass main airflow tube. Borosilicate glass electrodes were filled with the sensillum lymph ringers solution ^65^ were inserted into the eye and individual sensilla as reference and recording electrodes, respectively. The ab5 sensilla were distinguished by their location on the antenna and their response to a small number of diagnostic odorants. No more than 3 sensilla were recorded per fly. Extracellular action potential recordings were determined with an EXT-02B amplifier (NPI) with a 5x gain headstage. Data were obtained and AC filtered (300-3,000 Hz) at 10 kHz with a PowerLab 4/35 digitizer and Lab Chart Pro v8 software (ADInstruments).

Odorant cartridges consisted of 14.6 cm Pasteur pipettes capped with 1 mL pipette tips. Prior to capping, 50 µl of odorant solution was pipetted onto a 13 mm antibiotic assay disc (Whatman) and inserted into each Pasteur pipette. Odorants were allowed to equilibrate for at least 30 min. To apply the odorants, air at 500 mL/min flowed for 500 ms through the cartridge, which was inserted into a hole in the main airflow tube. Odor delivery was controlled by LabChart, which directed the opening of a valve linked to a ValveBank 4 controller (Automate Scientific). Cartridges were used up to 4 times, with at least 10 min recovery between trials on different sensilla. A 10 second trace was collected, including a 1 second baseline prior to odor application. Each sensillum was tested with multiple odorants with at least 10 seconds between odor applications.

For recordings in Fig. 4, an ab5 sensillum was first identified with the standard recording setup described above. Then, paired pulses and trains of geranyl acetate stimuli were applied to the sensillum with a modified delivery system. For this, a 1000 mL Erlenmeyer flask was filled with 200 mL of 0.01% geranyl acetate. The flask was capped using a rubber stopper with 2 openings. Through one opening, an input line was inserted. The other opening contained the output line that connected to the main airflow tube. A Lee valve upstream of the input line was used to divert a stream of 500 ml/min air through the geranyl acetate flask and into the main airflow tube. LabChart was programmed to change odor delivery systems and to apply the paired pulse and spike train stimuli. The three paired pulse stimuli were obtained were applied in a continuous recording by applying 500 ms of geranyl acetate at 1, 11, 26, 29, 44, and 45.5 seconds. Spike trains were obtained by applying 200 ms of geranyl acetate every second for five seconds.

Action potentials were detected offline using LabChart Spike Histogram software. Spikes from ab5A and ab5B neurons were summed due to their similar amplitudes ^66^. Action potentials were counted over a 500 ms window, 100 ms after stimulus onset to account for the line delay. Solvent corrected odor responses graphed in Fig. 3A, 3B, and 5 were determined as the number of spikes induced by the odorant after subtracting the number of spikes produced by stimulating with paraffin oil alone. In Fig. 4D, solvent corrected responses were used to calculate paired-pulse ratios as Response_2_/Response_1_. In Fig. 4F, solvent corrected responses were normalized by dividing each response by the mean response to the first pulse of geranyl acetate for that genotype. For the peristimulus time histograms (PSTHs) in Fig. 3C, spikes were identified in LabChart with the Spike Histogram feature, and spike timing data were exported to Igor Pro 8.04 (WaveMetrics) where responses were averaged in 50 ms bins to generate PSTHs for each sensillum. PSTHs were then averaged to obtain the mean ± SEM PSTHs.

## Acknowledgements

We thank John Carlson and the Bloomington Drosophila Stock Center (NIH P40OD018537) for fly lines used in this study. The pBGRY plasmid and *LexA::VP16* entry clone were gifts from John Carlson and Tong-Wey Koh. The pU6-BbsI-chiRNA plasmid was a gift from Melissa Harrison, Kate O’Conor-Giles, and Jill Wildonger. The pHD-DsRed plasmid was gift from Kate O’Connor-Giles. We thank the Drosophila Genomics Resource Center (NIH grant P40OD010949) for cDNA clones. We appreciate assistance from Olivia Durham and Alina Vulpe for cloning constructs for the *Cyp308a1-LexA* line. We thank Anastasios Tzingounis for comments on the manuscript. Research in K.M.’s laboratory was supported by NIH awards R03DC015629 and R35GM133209. D.A. was supported by NSF REU Award to the University of Connecticut 1852486.

## Contributions

K.M. conceived the project. RNASeq was performed and analyzed by S.B., P.M., and K.M. The qRT-PCR experiments were performed by S.B. Histology was performed by S.B., H.D., R.S., M.N., and D.A. The mutant and transgenic flies were generated by S.B., R.S., and M.N. All electrophysiological recordings were performed by P.M. The manuscript was written by K.M. with input from all authors.

## Figure Legends

**Supplementary Fig. S1:**
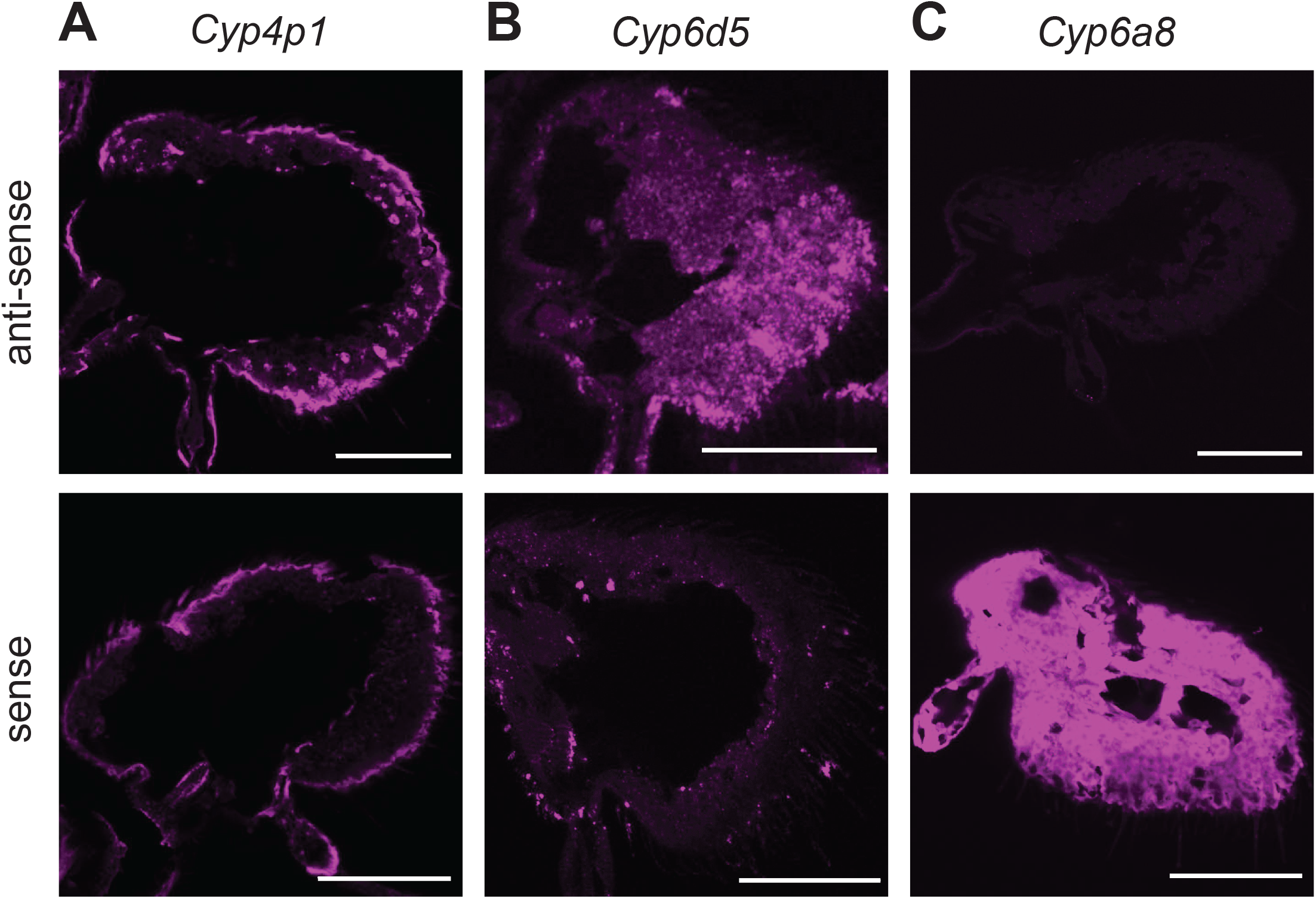
FISH can be used to localize *Cyp4p1* and *Cyp6d5*, but not *Cyp6a8*. (A) Antennal sections from wild-type flies exposed to 5% geranyl acetate for two hours show cellular labeling with an anti-sense *Cyp4p1* FISH probe, but not a sense control probe. Non-specific cuticular labeling on the edge of the antenna is also observed. (B) Similar to (A), but for *Cyp6d5*. (C) A *Cyp6a8* anti-sense FISH probe does not label antennal sections from wild-type flies exposed to geranyl acetate, but non-specific staining is seen in sense probe controls. Scale bars are 50 µm in each image.

**Supplementary Table S1:** CYP expression measured with RNASeq. The first 10 columns (A-J) list every CYP expressed in the DMSO exposed antennae with >5 RPM. The expression in each of three samples in each of three conditions (DMSO exposed, geranyl acetate exposed, and ethyl lactate exposed) is listed in RPM. Columns M-O list the mean expression in each condition. Columns R-T list the log_2_(RPM) in each condition. Columns W-Y list the results of the EdgeR ExactTest for differential gene expression for each comparison. The FDR rates are shown.

